# Integrating spatial transcriptomics data across different conditions, technologies, and developmental stages

**DOI:** 10.1101/2022.12.26.521888

**Authors:** Xiang Zhou, Kangning Dong, Shihua Zhang

## Abstract

With the rapid generation of spatial transcriptomics (ST) data, integrative analysis of multiple ST datasets from different conditions, technologies, and developmental stages is becoming increasingly important. However, identifying shared and specific spatial domains across ST datasets of multiple slices remains challenging. To this end, we develop a graph attention neural network STAligner for integrating and aligning ST datasets, enabling spatially-aware data integration, simultaneous spatial domain identification, and downstream comparative analysis. We apply STAligner to the integrative analysis of ST datasets of the human cortex slices from different samples, the mouse olfactory bulb slices generated by two profiling technologies, the mouse hippocampus tissue slices under normal and Alzheimer’s disease conditions, and the spatiotemporal atlases of mouse organogenesis. STAligner efficiently captures the shared tissue structures across different slices, the disease-related substructures, and the dynamical changes during mouse embryonic development. Additionally, the shared spatial domain and nearest neighbor pairs identified by STAligner can be further considered as corresponding pairs to guide the three-dimensional reconstruction of consecutive slices, achieving more accurate local structure-guided registration results than the existing method.

## Introduction

In multicellular organisms, cells are deeply influenced by their tissue environment. Knowledge of the spatial position of cells plays a critical role in understanding why cells from different locations have distinctive structures and functions^1^. Recent ST technologies provide new ways to measure mRNA expression and corresponding spatial coordinates in tissues^2^. Various ST technologies have been developed with different spatial resolutions. For example, the spatial barcoding-based method 10x Visium achieves a spatial resolution of 55μm with each capture site (i.e., spot) containing approximately 1-10 cells^3^. Slide-seq improves spatial resolution to near-cellular level (10μm)^4,5^, and Stereo-seq further reaches sub-cellular resolution (0.22μm)^6^. Indeed, these breakthrough technologies have been widely used to explore the biological architecture of tissues^4,5,7^, the regulatory mechanism of complex disease^8^, and the development process of organogenesis^6^. In particular, computational methods have been successfully applied to ST data of a single condition, technology, or developmental stage to characterize the tissue structure^9–12^, identify spatially variable genes^13,14^, and decipher spatial cell-cell communication patterns^15^ and so on. However, integrated analysis of ST datasets produced by different conditions, technologies, or developmental stages remains challenging.

Identifying the shared and specific spatial domains (defined as regions with similar spatial expression patterns) is one of the fundamental tasks for the integrative analysis of multiple ST datasets. Several methods have been proposed to address this issue in the single-slice ST dataset by incorporating spatial information and gene expression profiles. For example, BayesSpace^11^ encourages neighboring spots to belong to the same spatial domain by imposing spatial neighbor structure into the prior of its Bayesian statistical model. STAGATE^12^ employs a deep learning model and leverages the graph attention mechanism to learn the spatial heterogeneity. However, integrative and comparative analysis of ST datasets poses a unique challenge, as the technology noises introduced by different sequencing platforms or distinct conditions may mask the actual biological signals. Moreover, different sampling locations of slices may lead to completely distinct spatial structures.

Compared to single-slice ST data analytics for deciphering spatial heterogeneity, integrative and comparative analysis of ST datasets from different individual samples, biological conditions, technological platforms, and developmental stages can provide more comprehensive characterizations of spatial tissue structures. For example, integrating slices from normal and diseased tissue can facilitate the identification of disease-specific spatial domains^16^. Joint analyses of ST slices across different embryonic stages can facilitate the identification of stage-specific spatial domains and common domains spread across the developmental processes (in the developing tissues)^6^. As the integrative analysis for single-cell RNA sequencing (scRNA-seq) data, ST slices from different studies also contain batch effects which may mask actual biological signals and hamper data integration and interpretation^9^. However, most current integration analyses either pool gene expression data across slices or employ existing batch correction methods developed for scRNA-seq data, such as Harmony^17^, and Seurat V3^18^, without considering the spatial coordinates. An unsupervised spatial embedding method, SEDR^9^, was combined with Harmony to embed ST datasets for batch correction. Since these two modules are optimized independently, their combined performance is limited. A recently developed method, PASTE^19^, integrates adjacent slices into a center slice to improve the clustering performance of spatial domains and achieve three-dimensional (3D) reconstruction. However, PASTE requires global and rigid similarity between adjacent tissue slices from the same tissue and is not suitable for diverse ST slices from different individuals, conditions, technologies, or developmental stages.

To this end, we develop a graph attention neural network STAligner for integrating and aligning ST datasets through generating batch-corrected spot embeddings, enabling spatially-aware data integration, simultaneous spatial domain identification, and downstream comparative analysis. Specifically, STAligner employs the graph attention auto-encoder to learn spot embeddings with gene expression and spatial location information and constructs the spot triplets across slices to guide the embedding alignment process. STAligner demonstrates superior performance on a diverse range of ST datasets including the human dorsolateral prefrontal cortex slices from multiple samples, the mouse olfactory bulb slices from two different technologies, the mouse brain sagittal anterior and posterior slices, the normal and Alzheimer’s disease slices, and the spatiotemporal atlases of mouse organogenesis. For the multiple stacked ST slices, STAligner employs the shared spatial domain and nearest neighbors to guide pairwise registration, and accurately perform coordinates alignment and 3D reconstruction.

## Results

### Overview of STAligner

STAligner takes the normalized gene expression and spatial coordinates of ST data from multiple tissue slices as input (**Fig. 1**). For each ST slice, STAligner constructs a spatial neighbor network between spots using the spatial coordinates. STAligner employs a graph attention auto-encoder neural network to extract spatially aware embeddings as used in STAGATE^12^, and constructs the spot triplets based on current embeddings to guide the integration and alignment process of different slices by attracting similar spots and discriminating dissimilar spots across slices. STAligner defines the triplet that contains anchor-positive and anchor-negative spot pairs. The former is defined as the mutual nearest neighbors (MNNs) with similar gene expressions but belong to two different slices, and the latter belongs to the same slice with different spatial positions and dissimilar expressions. STAligner introduces the triplet loss to update the spot embedding to reduce the distance from the anchor spot to the positive one, and increase the distance from the anchor spot to the negative one. The graph attention auto-encoder training and triplet construction are optimized iteratively until batch-corrected embeddings are generated.

**Fig. 1.**
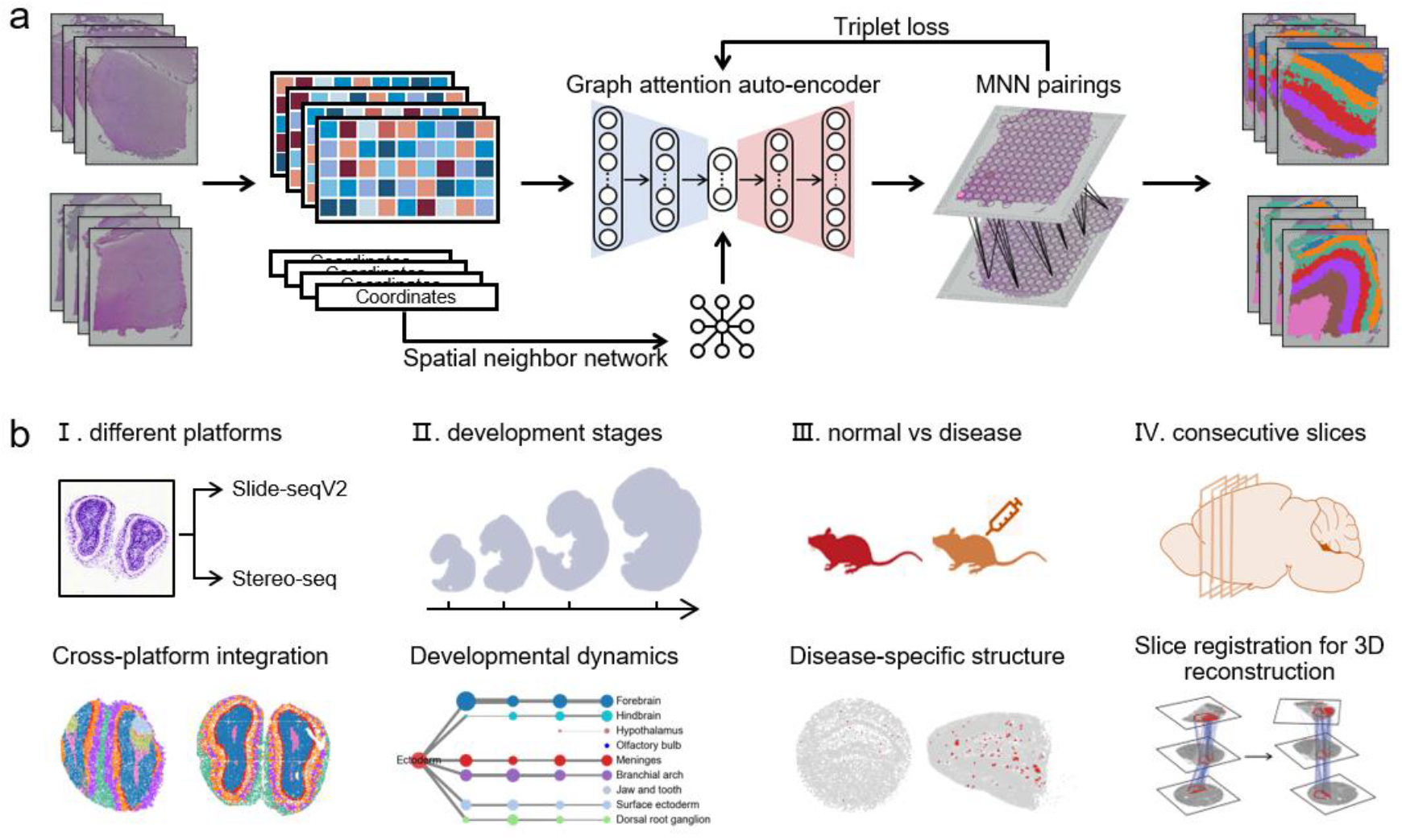
Overview of STAligner. **a**. STAligner first normalizes the expression profiles for all spots and constructs a spatial neighbor network using the spatial coordinates. STAligner further employs a graph attention auto-encoder neural network to extract spatially aware embedding, and constructs the spot triplets based on current embeddings to guide the alignment process by attracting similar spots and discriminating dissimilar spots across slices. STAligner introduces the triplet loss to update the spot embedding to reduce the distance from the anchor to positive spot, and increase the distance from the anchor to negative spot. The triplet construction and auto-encoder training are optimized iteratively until batch-corrected embeddings are generated. **b**. STAligner can be applied to integrate ST datasets to achieve alignment and simultaneous identification of spatial domains from different biological samples in (**a**), technological platforms (I), developmental (embryonic) stages (II), disease conditions (III) and consecutive slices of a tissue for 3D slice alignment (IV).

With the aligned embeddings, STAligner can be applied to integrate ST datasets to achieve simultaneous identification of spatial domains from diverse tissue slices. We applied STAligner to five scenarios to demonstrate its effectiveness including integrating ST slices across individual samples, technological platforms, embryonic stages, disease conditions, and stacked slice registration for 3D reconstruction (**Fig. 1**).

### STAligner improves the integration of multiple slices on the human dorsolateral prefrontal cortex dataset

To quantitatively evaluate the integration performance of STAligner, we applied it onto the human dorsolateral prefrontal cortex (DLPFC) ST dataset^7^ generated by 10x Visium. It consists of 12 tissue slices from three adult samples with four adjacent slices for each. The original study has manually annotated six neocortical layers from layer 1 to layer 6 and the white matter (WM) (**Fig. 2a**). By taking the manual annotations as ground truth, we evaluate the performance of STAligner on simultaneous identification of spatial domains and compare it with that of the non-spatial integration method Harmony and two recently developed spatial integration approaches SEDR and PASTE in terms of the Adjusted rand index (ARI) (see **Methods**).

**Fig. 2.**
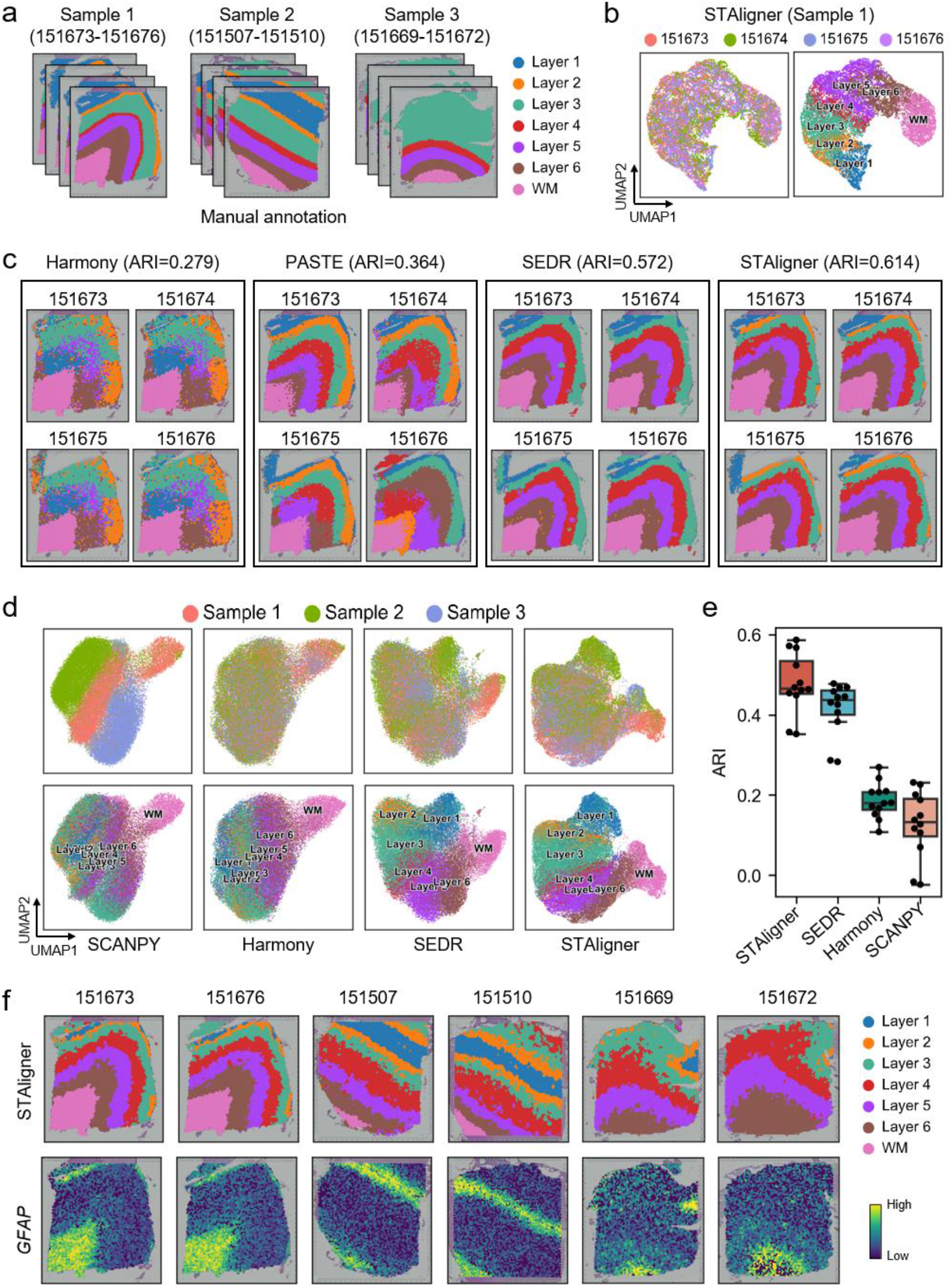
STAligner outperforms benchmarking methods on the human dorsolateral prefrontal cortex (DLPFC) slices from the same and different samples. **a.**Manual annotations of cortical layers and white matter (WM) on the slices of three DLPFC samples. **b.**UMAP plots of STAligner embeddings colored by slices (left) and cortical layers (right) for integrating and aligning four slices of sample 1. **c.**Aligned spatial domain identified simultaneously by integrating four slices of sample 1 via Harmony, PASTE, SEDR and STAligner respectively. **d.**UMAP plots of embeddings colored by slices (top) and cortical layers (bottom) by integrating and aligning 12 slices of all three samples via SCANPY, Harmony, SEDR and STAligner respectively. **e.**Boxplot of clustering accuracy of spatial domains identified simultaneous by the four methods for all 12 slices in terms of ARI scores. **f.**Spatial domains identified simultaneously by STAligner (top) and spatial expression of marker gene *GFAP* for layer 1 (bottom) in six slices respectively.

We first evaluate the performance on adjacent slices of a single sample and observe that both STAligner and SEDR can effectively integrate the four slices with well-matched low-dimensional embeddings (**Fig. 2b** and **Supplementary Fig. S1a**). The common cortical layers across slices are evenly mixed, and different layers are clearly separated and ordered with spatial structure from layer 1 to layer 6 and WM. While Harmony cannot separate the cortical layers clearly, even its embeddings of the four slices are mixed. PASTE can capture the layer structures but the slices are not evenly mixed since it does not perform embedding alignment explicitly. STAligner achieved the highest ARI scores among all methods for the three samples respectively. For example, STAligner accurately captured the layer boundaries and obtained higher clustering accuracy (ARI=0.61) than that of Harmony (ARI=0.28), PASTE (ARI=0.36), SEDR (ARI=0.57) for sample 1 (slices 151673-151676, **Fig. 2c**). We can find that the layer structure identified by the non-spatial approach Harmony suffers from serious noisy boundary issues and could only identify the layer 3 and WM structure correctly. For comparison, the spatially aware embedding methods could more accurately capture tissue architecture heterogeneity and the obtained domain boundary is significantly smoother. Similar conclusions can also be found for the other two samples (151507-151510 and 151669-151672), indicating the importance of considering spatial information for integration of multiple ST slices even from the same tissue sample (**Supplementary Fig. S1b**).

We further compare the performance on more challenging setting by integrating all 12 slices from the three samples. The UMAP plot of unaligned embeddings produced by SCANPY showed that the three samples exhibited substantial batch effects with common cortical layers from different samples separated (**Fig. 2d**). The embeddings of Harmony and SEDR were overcorrected with disordered layer structure, while STAligner can evenly mix the spots from the shared layers across samples and make the layers organized as their spatial locations. This conclusion can be further confirmed by quantitative measure of batch-corrected embedding (**Supplementary Fig. S2a**), where STAligner achieved comparable divergence score but much higher silhouette coefficient than competing methods. Moreover, the previously reported layer-marker genes^7^, such as *FABP7* (layer 1), *HPCAL1* (layer 2), *CARTPT* (layer 3), *PCP4* (layer 5), *KRT17* (layer 6), *MBP* (WM), showed differential expressions among the identified layers (**Supplementary Fig. S2b**). STAligner demonstrated more consistent spatial domains across samples (**Supplementary Fig. S2c**), and higher median ARI score of 0.47, compared with that of Harmony (median ARI=0.19) and SEDR (median ARI=0.43) (**Fig. 2e**). In addition, STAligner recognized the structure of layer 1 and layer 2 at the top right part of layer 3 in sample 2, which were missed when analyzing this sample alone and can be verified by the expression of marker gene *GFAP* (**Fig. 2f**), demonstrating STAligner could clearly depict the shared tissue structures across different samples.

Some ST technologies may be limited by the size of the captured area and cannot cover a whole organ of interest for some cases. As a solution, the tissue sample is dissected into multiple slices. We applied STAligner to a mouse brain dataset where the tissue was divided into the sagittal anterior and posterior slices (**Supplementary Fig. S3a**). STAligner clearly captured the known brain tissue structures of olfactory bulb (domains 3, 17, 37), cortex (domains 6, 9, 10, 16, 20, 33, 35), stratium (domains 2, 12, 34), thalamus (domains 4, 24, 27, 30), hypothalamus (domains 5, 7, 28), hippocampus (domains 14, 19, 31, 36), cerebellum (domains 8, 11, 22, 32), midbrain (domains 18, 23), hindbrain (domains 1, 13, 25, 29, 39), and corpus callosum (domains 14, 21) (**Supplementary Fig. S3b**). Particularly, STAligner accurately recognized three small structures in the hippocampus region, with the domain 31 denoting dentate gyrus structure (DG), the domain 14 denoting CA1, and the domain 36 denoting CA3 (**Supplementary Fig. S3c**). We also found that the tissue structures flanking the shared border of the anterior and posterior slice were accurately identified as the same domains, such as the structures CA1 and CA3, and the superficial and deep cortex layer (domains 10, 16, 20), which were also clearly characterized by some marker genes like *Fibcd1* in CA1, *Cabp7* in CA3^20^, *Lamp5* in superficial cortex layer^7^, and *Fezf2* in deep cortex layer^21^. Thus, STAligner could well stitch multiple tissue slices into a large one for deciphering its global structure.

### STAligner identifies common and specific tissue structures across two mouse olfactory bulb ST slices profiled by different technological platforms

Here we applied STAligner to integrate two mouse olfactory bulb ST slices profiled by Slide-seqV2^5^ and Stereo-seq platforms^6,9^ respectively (**Fig. 3a**) with clear tissue structures (**Fig. 3b**) but substantial batch effects (**Supplementary Fig. S4**). Harmony followed with Louvain clustering failed to recover the laminar organization, and SEDR found some intermixed spatial clusters with unclear tissue structures (**Fig. 3c**). By contrast, STAligner can well decipher the known tissue structures according to the annotated laminar structure in the DAPI-stained image^9^ and Allen Mouse Brain Atlas annotation (**Fig. 3b**) from both slices, indicating these two platforms indeed capture most of the key tissue structures. The six shared spatial domains by the two slices were spatially ordered from inner layers, middle layers, to outer layers of mouse olfactory bulb, including the rostral migratory stream (RMS), granule cell layer (GCL), mitral cell layer (MCL), external plexiform layer (EPL), glomerular layer (GL) and olfactory nerve layer (ONL) (**Fig. 3c-e**). Moreover, STAligner identified two slice-specific spatial structures, the accessory olfactory bulb (AOB) and the granular layer of the accessory olfactory bulb (AOBgr) in the Slide-seqV2 ST slice, which may be caused by different sampling locations (**Fig. 3c-e**). The UMAP visualization of STAligner embeddings clearly demonstrated that the slice-shared laminar structures were well mixed and the slice-specific structures were clearly separated. However, Harmony and SEDR cannot recover these two structures and reveal such differences between these two ST slices (**Supplementary Fig. S4**). Furthermore, these spatial domains identified by STAligner were clearly characterized by known marker genes (**Fig. 3e**). For example, the expression of *Fxyd6^22^* displayed distinct spatial distribution on the recognized AOB domain, and the expression of the granular cell marker *Atp2b4^23^* showed distinct spatial distribution on the recognized AOBgr domain. The narrow loop structure of MCL was enriched in the spatial expression of mitral cell marker *Gabral^24^*. These results demonstrate that STAligner can integrate slices with different tissue structure compositions and avoid overcorrection of the slice-specific spatial domains.

**Fig. 3.**
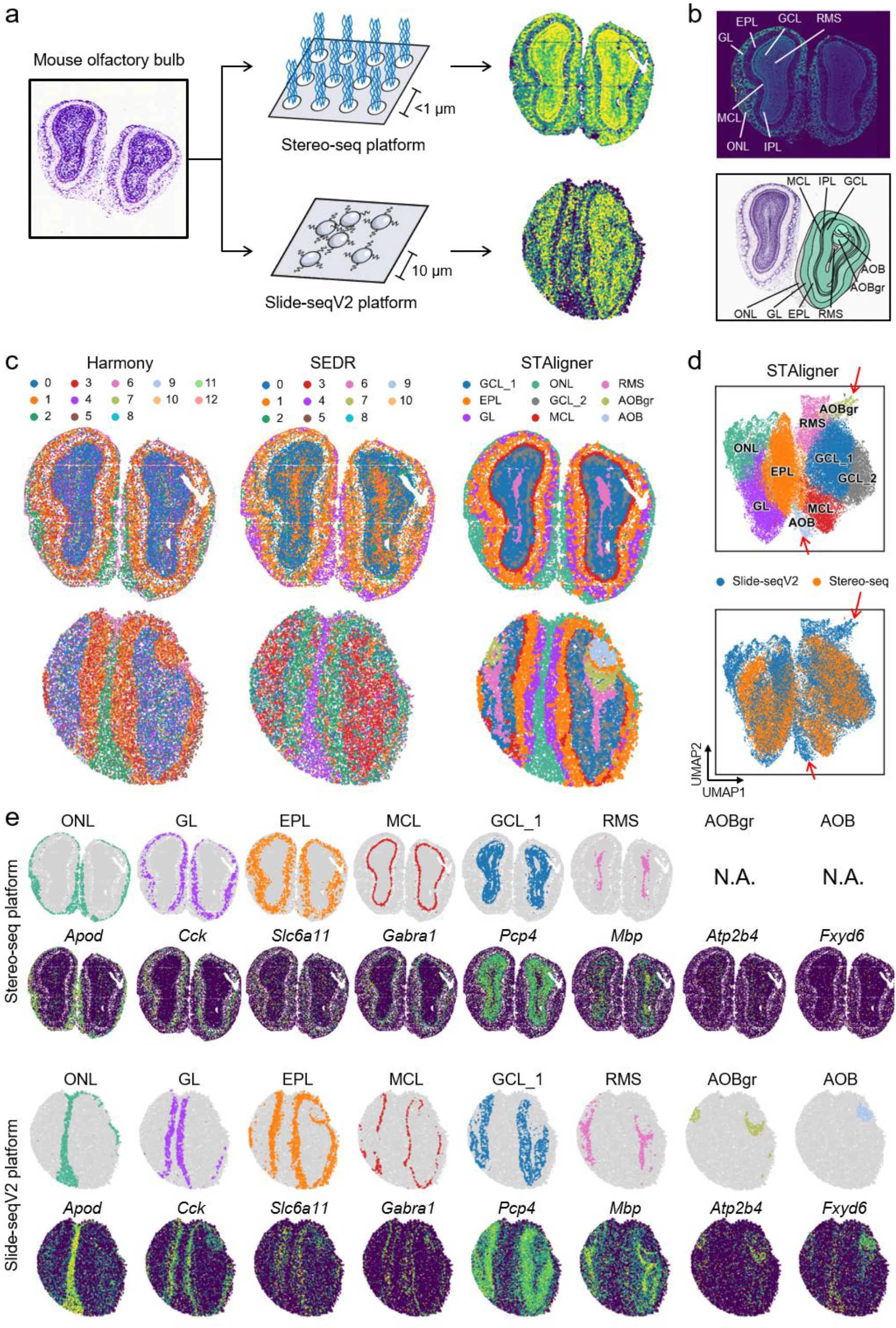
STAligner accurately identifies common and unique tissue structures in the mouse olfactory bulb data across two different sequencing platforms (i.e., Stereo-seq and Slide-seqV2). **a.**Two mouse olfactory bulb datasets produced by Stereo-seq and Slide-seqV2, respectively. **b.**Laminar organization of mouse olfactory bulb annotated by DAPI-stained image (top) and the Allen Reference Atlas (bottom). **c.**Visualization of spatial domains identified simultaneously by integrating and aligning the two ST slices profiled via Harmony, SEDR, and STAligner respectively. **d.**UMAP plots of STAligner embeddings colored by spatial domain (top) and slices (bottom). **e.**Spatial domains identified simultaneously by STAligner and the corresponding marker genes in the slice of Stereo-seq (top) and Slide-seqV2 (bottom) respectively.

### STAligner reveals detailed developmental dynamics of anatomical structures during mouse organogenesis

We applied STAligner to integrate four mouse embryo slices sampled at the time stages of E9.5, E10.5, E11.5 and E12.5 profiled by Stereo-seq^6^ to investigate the spatiotemporal heterogeneity of tissue structures during mouse organogenesis. Although the four slices have very different sizes and there are obvious batch effects, STAligner can effectively integrate the four slices in the embedding space and align the spatial domains which showed very consistent localization spatially across the four embryonic stages (**Fig. 4a**, **Supplementary Fig. S5a**). Each domain showed clear tissue structure and can be well annotated based on the anatomic location using eHistology Kaufman Annotations^25^ (**Fig. 4a**). To reconstruct the developmental trajectory of each tissue structure, we built connections between the same spatial domains in successive stages and generated an acyclic directed graph to reflect the ancestor-descendant relationships (**Fig. 4b**). According to the origination from the three germ layers during gastrulation, the annotated organs and tissues can be summarized into three parts, i.e., ectoderm, mesoderm, endoderm. Notably, the tissues originated from different germ layers also shown clear co-localization patterns based on the UMAP visualization (**Supplementary Fig. S5a**). More importantly, the annotated tissues across the four stages were well characterized with some known marker genes^6^, such as *Sox2* for the forebrain, *Ina* for the hindbrain, *Tbr1* for the olfactory bulb, *Six6* for the hypothalamus, *Myl7* for the heart, *Trf* for the liver, *Pax1* for the cartilage primordium, and *Myog* for the muscle (**Fig. 4c**, **Supplementary Fig. S5b**).

**Fig. 4.**
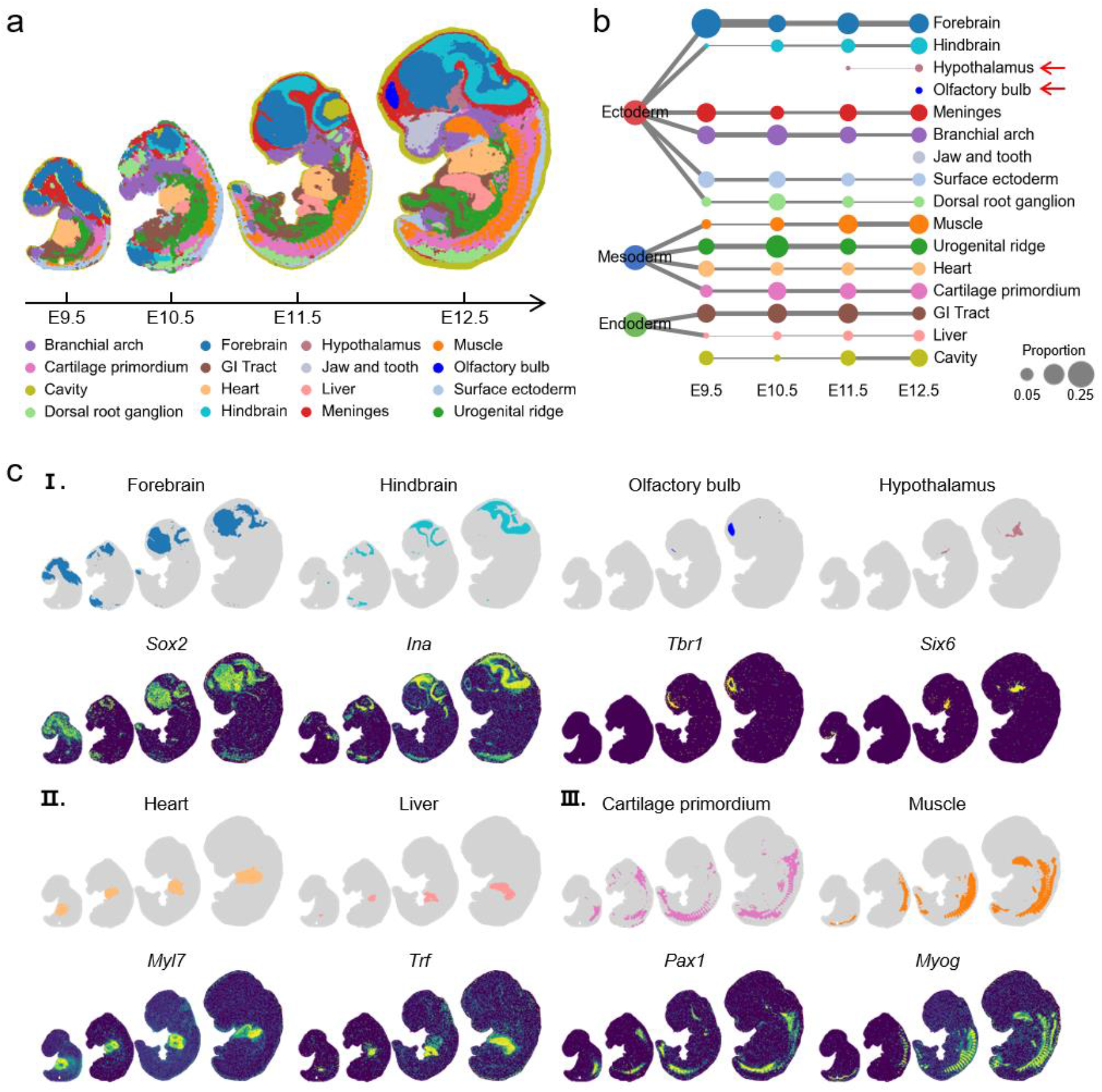
STAligner reveals developmental dynamics of anatomical tissue substructures (spatial domains) in the early mouse embryo data. **a.**Aligned spatial domains characterized simultaneously by integrating and aligning four ST data of four embryonic slices (E9.5 to E12.5) via STAligner. **b.**Directed acyclic graph showing the developmental dynamics of tissue substructures. Each row corresponds to one of the tissue substructures (spatial domains) shown in (**a**). **c.**Spatial tissue substructures relating to brain (I), heart, liver (II), cartilage primordium and muscle (III) and their corresponding marker genes (bottom).

All the major tissues and organs (e.g., heart, liver and brain) were well detected and consistent with well-known anatomical structure of the organ (**Fig. 4b** and **c**). Particularly, the heart was accurately identified with nearly unchanged proportions over the course of the developmental period, confirmed the fact that the heart is the first organ formed and functioned in the early embryo^26^. The liver occupies a small area at the earliest stage E9.5 and rapidly increases size in the following stages. Likewise, we observed prominent increase in the proportions of hindbrain and muscle, indicating the proliferation of these tissue structures during the four stages of organogenesis. In addition, STAligner can also delineate tissues with special shapes, such as the meninges with a continuous closed loop at late embryonic stages. STAligner can also detect small (proportion<5%) structures, such as hypothalamus and olfactory bulb, at later developmental stages (after E11.5), which are supported by known histological evidence. The hypothalamic neurogenesis starts at E11.5 and reaches the peak at E12.5^27^. The development of olfactory bulb initiated at E10.5 and cannot be detected macroscopically until E12.5^28,29^. These results demonstrate that STAligner can well integrate the ST slices of different developmental stages, facilitate the cross-stage comparison and deepen our understanding on the developmental dynamics of the anatomical structure changes during mouse organogenesis.

### STAligner well deciphers the amyloid-beta (Aβ) plaque-related cluster by integrating two ST slices of normal and Alzheimer’s disease mouse hippocampus tissue

Here we applied STAligner to two ST slices from normal^5^ and Alzheimer’s disease (AD)^8^ mouse hippocampus tissue profiled by Slide-seqV2, and identified 16 distinct spatial domains. Most spatial domains revealed clear spatial patterns and corresponded to anatomical structures of hippocampus in the Allen Reference Atlas (**Fig. 5a, b**, **Supplementary Fig. S6a**). The UMAP visualization of STAligner embeddings clearly showed that the common domains from the two slices were highly overlapping and slice-specific domains were well separated (**Fig. 5c**). STAligner could well capture shared tissue structures between these two conditions. For example, the domains 4, 7, and 11, construct a clear “cord-arrow-like” structure in the hippocampal region in both the two slices (**Fig. 5d, Supplementary Fig. S6a**). According to the Allen Reference Atlas, the domain 4 (“arrow-like”) corresponds to the dentate gyrus (DG) region, and the domains 7 and 11 (“cord-like”) correspond to the structure CA1 and CA3 of Ammon’s horn, respectively, which were well supported by some known gene markers. For example, *Lrrtm4* regulating excitatory synapse development and AMPA receptor trafficking in dentate gyrus granule cells was clearly expressed in the DG field^30^.

**Fig. 5.**
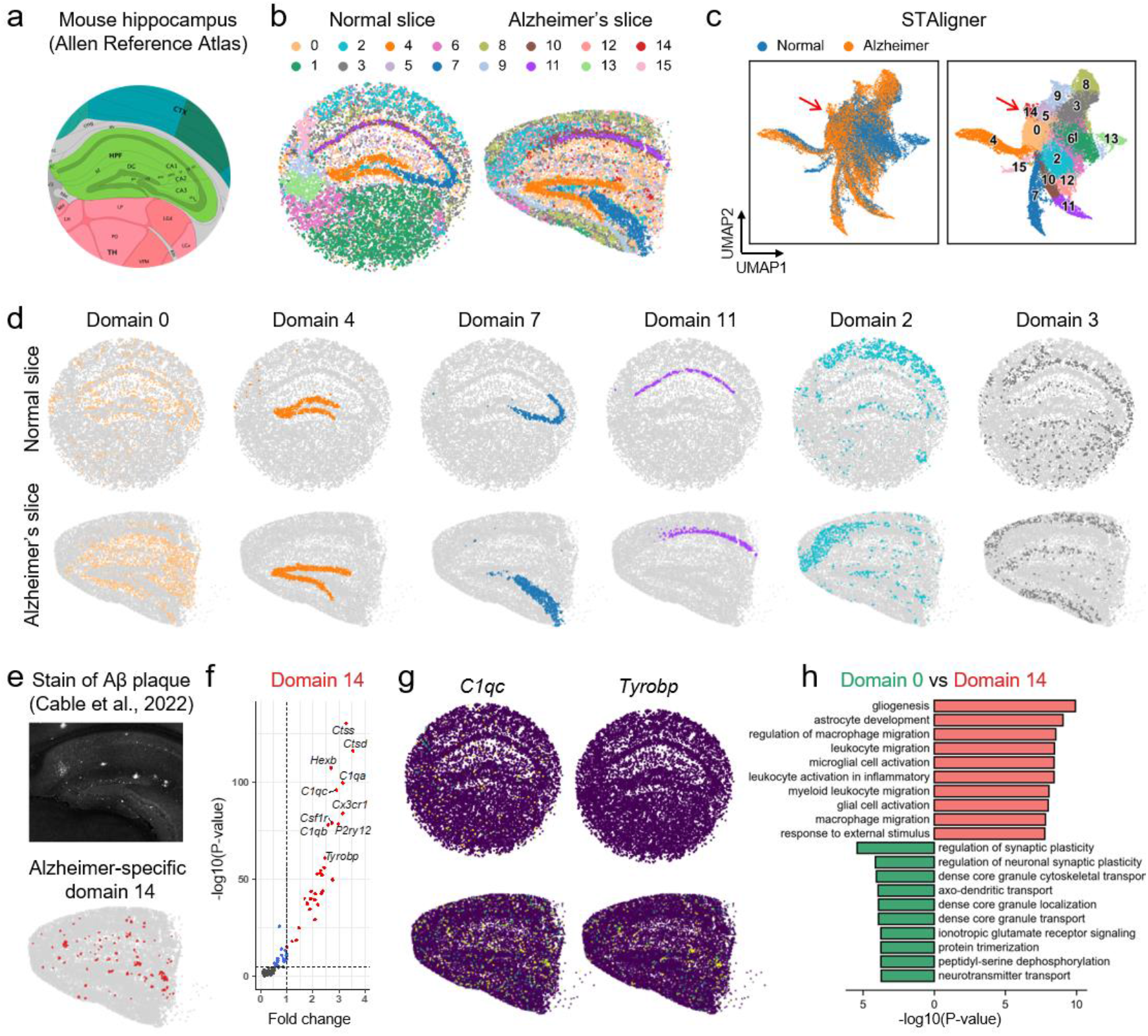
STAligner could well integrate and align two mouse hippocampus ST slices of normal and Alzheimer’s disease conditions. **a.** Laminar organization of mouse hippocampus annotated by the Allen Reference Atlas. **b.** Aligned spatial domains characterized simultaneously by STAligner between the normal and Alzheimer’s mouse hippocampus datasets produced by Slide-seqV2. **c.** UMAP plots of STAligner embeddings colored by condition types and spatial domains. **d.** Visualization of spatial domains identified simultaneously between the two datasets by STAligner. **e.** Spatial domain 14 was dominantly found in the Alzheimer’s slice (bottom) and co-localized with the Antibody stain of Aβ plaque in the adjacent hippocampus slice (top) (adapted from Cable et al., 2022). **f.** Volcano plot of DEGs between domain 14 and domain 0. **g.**Spatial expression maps of two marker genes *C1qc* and *Tyrobp* for spatial domain 14 in the normal and Alzheimer’s slice. **h.** GO enrichment terms of the DEGs.

**Fig. 6.**
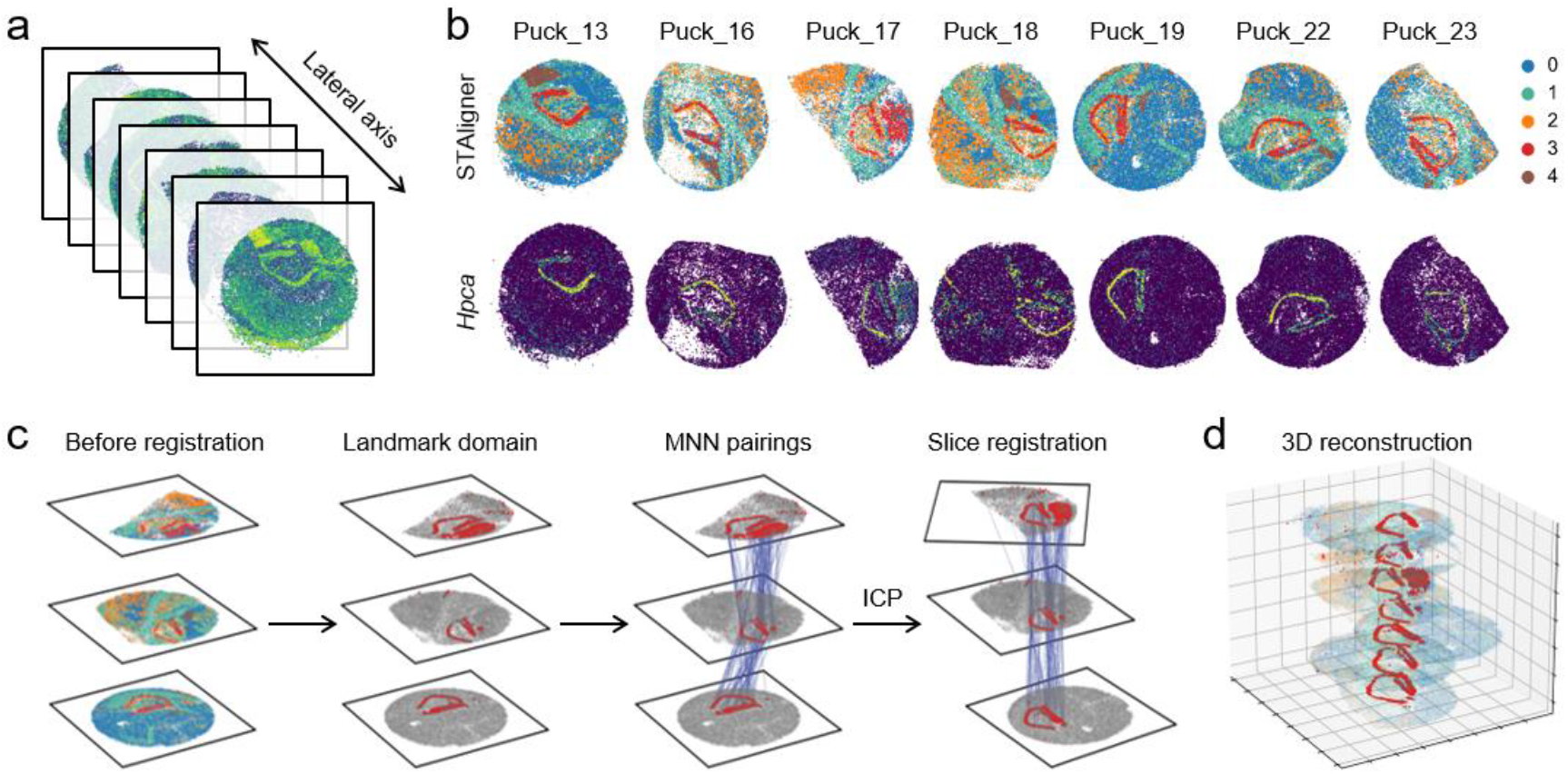
STAligner enables stacked 3D alignment of adjacent mouse hippocampus slices. **a.**Seven mouse hippocampus slices produced by Slide-seq. Spots in slice are colored by the number of transcripts. **b.**Spatial domains identified simultaneously by integrating and aligning seven consecutive slices (top) via by STAligner and the expression of marker gene *Hpca* for domain 3 (bottom). **c.**The workflow of downstream registration including landmark domain extraction shared by adjacent slices, MNN pair search between adjacent slices as anchor pairs, and pairwise registration using the ICP algorithm. **d.**Well registered slices with the hippocampus structure highlighted.

By computing the composition of each domain belonging to AD or normal slice (**Supplementary Fig. S6b**), we found that domains 0 and 14 located at the central hippocampal region (except for the “cord-arrow-like” structure), but showed clear different compositions between slices, where domain 14 was more enriched in the AD slice. From the UMAP visualization, we can observe that domain 14 is localized in close proximity to domain 0 (**Fig. 5c**). Importantly, domain 14 was highly co-localized with the amyloid deposition in the Aβ plaque staining image (**Fig. 5e**). These observations suggested that domain 14 was AD-associated. Moreover, the highly expressed genes (significantly upregulated) in domain 14 compared with domain 0 included microglia marker genes and some of them were implicated to the AD risk (**Fig. 5f**). For instance, *Csf1r* is a key growth factor receptor for microglia and plays a crucial role for its homeostasis and development^31^; and *Tyrobp* was found to be markedly upregulated in Aβ-associated brain regions^32^ and could enhance phagocytic activity of microglia and clear Aβ deposits^33^. Early study demonstrated that *C1qa, C1qb*, and *C1qc* were strongly expressed for Aβ-associated microglia and could mediate synapse loss in AD^34^. We can also find that the spatial expression distribution of DEGs (*C1qc* and *Tyrobp*) localized near the position of Aβ accumulation, illustrating the potential effect of Aβ accumulation on gene expression (**Fig. 5g**). These DEGs were involved in many gene ontology (GO) terms such as gliogenesis, astrocyte development, microglial cell activation (**Fig. 5h**). These results demonstrate STAligner could preserve true biological variations while remove batch effects, and untangle the dysregulated cellular network surrounding the Aβ deposition of AD.

### STAligner enables stacked 3D tissue reconstruction of consecutive mouse hippocampus slices

For the same tissue, multiple consecutive adjacent tissue slices need to be manually registered by rigidly rotation and translation for 3D tissue reconstruction. Here, we demonstrate the capability of automatic registration of seven adjacent mouse hippocampus slices generated by Slide-seqV1^4^ (**Fig. 6a**). It is challenging to integrate and register these slices due to the low transcript detection sensitivity of Slide-seqV1 and only partial overlap among these adjacent slices. STAligner could successfully integrate the seven mouse hippocampus slices, and recognize the shared structure of corpus callosum (domain 1), Ammon’s horn and the dentate gyrus (domain 3), and third ventricle (domain 4), which demonstrated high consistency with the hippocampus annotations in the Allen Reference Atlas (**Fig. 5a** and **Fig. 6b**). The UMAP visualization of the STAligner embeddings demonstrated that the spots across slices were mixed well while the spatial domains were clearly separated (**Supplementary Fig. S7a**) with clear expression pattern (**Supplementary Fig. S7b**). These tissue structures were further confirmed by the higher expressions of oligodendrocytes marker *Plp1^35^* (corpus callosum), *Hpca^36^*(Ammon’s horn and the dentate gyrus), and choroid plexus marker *Ttr^37^* (third ventricle). Due to the difference between sampling position in the experimental procedures, the tissue in Puck_16-18, 22-23 were incomplete, resulting in the differences of the shape and distribution of the tissue structures across the seven tissue slices. However, embedding alignment does not mean geometrical registration. According to the orientation and position of domains shared across slices (domain 3), we can observe that the adjacent slices are clearly misaligned (**Fig. 6b** bottom panel).

Intuitively, using a common domain with asymmetrical shape and enough overlapping areas could make the further registration process robust against noise. We used the domain 3 as a landmark domain, which has a clear “cord-arrow-like” structure. The MNNs in this domain identified by STAligner were selected as anchor pairs to guide the registration process using the ICP algorithm (**Fig. 6c**). All adjacent slices are aligned sequentially with translation and rotation along the z axis. From the stacked 3D tissue with registered coordinates, we can observe that the orientation and position of the “cord-arrow-like” structure are matched accordingly (**Fig. 6d**), resulting in a clear visual improvement of 3D spatial patterns. By contrast, PASTE that ignores the local spatial structures, cannot correctly align the domain 3 across adjacent slices (**Supplementary Fig. S8**). These results illustrate that the MNNs and spatial domains generated by STAligner could help to guide the coordinate registration of adjacent slices and achieve promising 3D tissue reconstruction for diverse tissue slices.

## Discussion

Integrative and comparative analyses of multiple ST slices are essential for a comprehensive understanding of the heterogeneous tissue structures across different samples, technical platforms, biological conditions and developmental stages. We develop a tool STAligner to integrate and align spots from multiple ST slices in a batch-corrected embedding manner. STAligner adopts the graph attention auto-encoder to learn spatially informed embeddings and the triplet learning to guide the embedding alignment process.

The rapid development of ST technologies results in higher spatial resolution, larger field of view, and larger number of spots. These large-scale datasets require the integration method to be memory and time-efficient. We recorded the computational time and GPU memory usage cost by STAlinger on three datasets in **Supplementary Fig. S9a**. When integrating the largest mouse hippocampus dataset with seven slices and more than 170k spots in total, STAligner is fast and only consumes 8.5 minutes. We can also find that the GPU memory usage remains constant when increasing the number of slices (**Supplementary Fig. S9b**), which makes STAligner suitable for analyzing large-scale ST datasets.

Currently, our 3D reconstruction strategy based on the ICP algorithm is limited to rigid transformation and cannot account for local flexible distortions. Future work is expected to develop spatial-domain guided flexible registration method for 3D reconstruction. The flexible transformation strategy can even be used to register slices from different samples while handling the anatomical differences across samples, and build a common reference of organs of different individuals and constructing ST atlases in the future^38^.

## Methods

### Data preprocessing

STAligner takes gene expression and spatial coordinate information of multiple ST slices as input. For each slice, we normalize the raw gene expressions according to library size and log-transformed using the SCANPY package^39^. Then, we select the top 5000 highly variable genes (HVGs). We take the intersection of the HVGs of all the input slices to ensure the spot features are shared across slices.

### Construction of the spatial neighbor graph

Based on the spatial coordinate information of all spots, we measure the similarities between spots by the Euclidean distance. Two spots are considered as neighbor if the distance between them is shorter than a pre-defined threshold *r*. Using the spatial neighbor relationship, we construct an undirected neighbor graph denoted as the adjacency matrix *A*, where *A*_ij_ = 1 if and only if there is an edge between spot *i* and spot *j*. *A* is a symmetric matrix with self-loops added for each spot. We can adjust *r* so that each spot has 4-15 nearest neighbors on average for diverse ST scenarios and platforms.

### Spatially aware spot triplet learning

STAligner consists of two parts including the graph attention autoencoder for spatially aware embedding learning and the spot triplet learning for batch correction (**Fig. 1**).

#### Graph attention auto-encoder

Graph attention auto-encoder^40^ is an unsupervised representation learning framework using both graph structure information and node attributes, which have been successfully adapted for deciphering spatial domains in single ST data in STAGATE^12^. It employs the attention mechanism to adaptively determine the influence of neighbor nodes on learning node representations. Specifically, the graph attention encoder takes the normalized gene expressions X and the corresponding graph structure as input, to get spot representations z. Let *L* be the number of layers in the encoder, *N* be the number of spots in the slices. The output representation of the *k*-th (*k*∈{1,2,..*, L* - 1}) encoder layer of spot *i*∈{1,2,.., *N}* is formulated as follows:

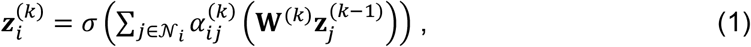

where 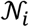 represents the neighbor set of spot *i* (including spot *i* itself), **W**^(*k*)^ is the trainable weight matrix, 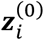 is the initial gene expression **x**_*i*_ of spot *i*. 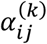 is set to make the relevance coefficients of spot ^i^’s neighbors comparable:

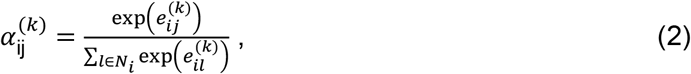

where 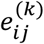 indicates the relevance of spot *i* to its neighboring spot *j*, and can be computed by:

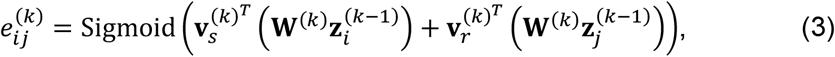

where 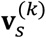 and 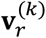 are the trainable weight parameters of the *k*-th layer, and Sigmoid denotes the *sigmoid* activation function. The *L*-th encoder layer does not use the attention mechanism and can be calculated by:

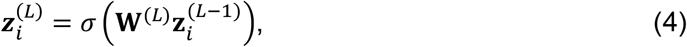

which is considered as the final embedding of spot *i*.

The number of layers of the decoder is the same as the encoder. To reconstruct expression features, the decoder reverses the process of learning latent embedding in the encoder. Using the output of the encoder as the input of the decoder, the *k*-th (*k* ∈ {2,..,*L* - 1,*L*}) decoder layer reconstructs the representation of spot *i* in layer *k* – 1 as follows:

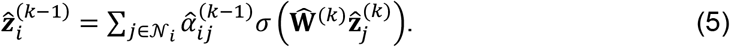

Similar to the encoder, the attention mechanism is not used in the last layer:

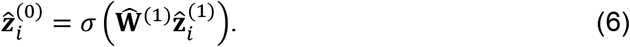

The output 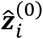 is considered as the reconstructed expressions. To avoid overfitting, STAligner sets 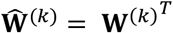 and 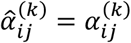 respectively.

The objective function of the graph attention auto-encoder is to maximize the similarity between input and reconstructed expressions, achieved by minimizing the mean square error loss:

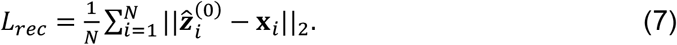

#### Spot triplet learning

We aim to enforce the encoder to learn a batch-corrected embedding for multiple ST data via the triplet learning, which has shown superior performance in diverse fields including single-cell RNA-seq dataset integration^41^. We define the spot triplet consisting of an anchor, a positive spot and a negative spot. It should be pointed out that, to avoid potential batch effects and dropouts, the triplets are defined in the embedding space instead of original gene expression space. The formed anchor-positive pair is used to overcome the batch effect while the anchor-negative pair guides the alignment in a discriminative way. We use the MNN pairs to define the anchor and positive spot which have similar gene expression but from two different slices. The negative spot is randomly sampled from the same slice as the anchor spot. Since the total number of spots is usually much larger than the *k*-nearest neighbors of the anchor spot, the probability of the sampled negative spot locating in the *k*-nearest neighbors of the anchor is very low. This negative sampling strategy can accelerate the model training especially in large-scale datasets. The triplet loss is used to minimize the distance between the anchor-positive pair and maximize the distance between the anchor-negative pair in the latent space z. It is formulated with a Euclidean distance function as follows,

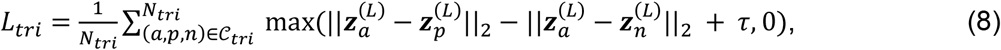

where *a, p* and *n* represent the indices of the anchor, positive, and negative spots respectively, *C_tri_* is the set of the identified spot triplets with size *N_tri_*, and *τ* (default = 1.0) is the margin used to enforce the distance between positive and negative pairs. Finally, the final objective is written as:

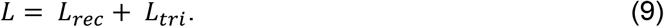

We perform spot triplet detection and graph attention auto-encoder training in an iterative manner to generate refined triplets and improved embeddings. We first detect spot triplets in the embedding space generated by the encoder from the last iteration, and then guide the embedding alignment process with the detected triplets. Here, the initial embeddings are generated without using the triplet loss.

### Landmark spatial domain guided consecutive slice registration and 3D tissue reconstruction

Registration of the spot coordinates of the adjacent slices is an important step for 3D tissue reconstruction. Typical registration process uses landmark points to guide the registration, including three steps: landmark point (key point) extraction, landmark point matching and the optimal transformation estimation. ICP algorithm^42,43^ is a typical method to achieve rigid transformation (i.e., translation and rotation) to register the coordinates from different coordinate systems to a common coordinate framework with given landmark point matching. However, it is challenging to directly apply the algorithm to adjacent slices due to partial overlapping and dropout noise. Instead of detecting the correspondences for each spot in the adjacent slices, we consider the MNN spots as landmark points which come from a selected landmark domain shared across slices. Here, the landmark domain should be asymmetrical and has enough overlapping areas to improve the robustness of the corresponding points. Using the coordinates of the MNN spots as the input of ICP algorithm, we could obtain a transformation matrix which can be further used to map all the spots of the target to reference slice. The registration process is performed in a sequential pairwise manner across all the adjacent ST slices.

### The overall architecture of STAligner

The encoder of STAligner consists of two layers with node dimensions set as 512 and 30 respectively, and the decoder is just the reverse of the encoder. The Adam optimizer^44^ is used to minimize the loss function with learning rate of 10^-4^. We set the weight decay as 10^-4^and the activation function as the exponential linear unit (ELU)^45^. We adopt the PyTorch Geometric library^46^ to implement the graph attention auto-encoder. We train the model in two steps: the graph auto-encoder model is first trained without the triplet loss (500 iterations by default), and generate the embeddings for triplet identification. Then, we alternate the model training and triplet refinement every 100 iterations until the fixed number of iterations reached (500 iterations by default).

### Clustering

The spatial domains were identified by clustering algorithm with the embeddings from STAligner as input. For datasets with prior knowledge on the number of domain label, the mclust clustering algorithm implemented by R package mclust (v 5.4.9) was used^47^. For dataset without prior knowledge, the Louvain algorithm implemented by SCANPY package (v 1.9.1) was employed. The adjusted Rand index (ARI) is used to quantify the agreement between the clustering result and the original annotation. Higher ARI score means better agreement between two grouping results while zero score means that the clustered labels are randomly guessed.

### Differential expression analysis and GO enrichment analysis

We used the Wilcoxon test implemented in the *FindMarkers(*) function of the R package Seurat (v4.1.0) to identify differentially expressed genes for the spatial domain in normal and disease slices with a cutoff of the adjusted *p* value (Benjamin-Hochberg correction) = 0.01. We performed the GO enrichment analysis for the domain-specific DE genes using the R package clusterProfiler (v4.2.2)^48^.

### Benchmarking integration methods

To benchmark the integration performance, we considered the following integration methods: Harmony^17^, SEDR^9^, and PASTE^19^. We also used SCANPY^39^ to produce uncorrected embeddings of the combined raw input data, where principal component analysis (PCA) was used to perform dimensionality reduction and no integration method was used. All parameters were kept at default values.

#### Harmony

Harmony has demonstrated its superior performance in a previous benchmarking study for scRNA-seq data integration^49^. According to the standard workflow, PCA was first performed on the combined expression matrix, and then the obtained PCA embeddings and batch information were considered as input of the *scanpy.external.pp.harmony_integrate()* function in the SCANPY package. Finally, the corrected PCA embeddings were obtained as output. The number of principal components (PCs) is set as 50 (default setting).

#### SEDR

SEDR is a spatial embedding representation method based on graph neural network with PCA embeddings as node features. In the original study, the authors fed the latent embeddings of SEDR to Harmony for batch correction. We used the demo code *run_UBC_DLPFC_data.py* in the online tutorial for unsupervised batch correction experiments. We empirically disabled the DEC loss function to obtain better performance. The PCs is set as 300, and the epoch is set as 300 (default setting).

#### PASTE

Using optimal transport that leverages both transcriptional similarity and spatial distances between spots, PASTE integrates multiple adjacent slices from the same sample in two alignment modes. The users can choose the mode by the biological problem of interest. The first mode finds an optimal probabilistic mapping between spots in two adjacent slices and uses the mapping to achieve spatial coordinate registration. By pairwise alignment between multiple pairs of adjacent slices, a stacked 3D spatial reconstructions can be obtained. We ran this mode on the seven mouse hippocampus slices (**Supplementary Fig. S8**). The second mode infers an enhanced ‘consensus’ slice by integrating multiple adjacent slices which can increase the power of downstream analysis. The low-dimensional embeddings in the integrated consensus slice can be used for further clustering analysis. We ran this mode on each DLPFC slice and concatenated the embeddings of each slice for evaluation. We ran PASTE according to the online tutorial *getting-started.ipynb* where the *pst.pairwise_align()* function and *pst.center_align()* function were used for the first and second mode, respectively. In the second mode, the size of low-dimensional representation is set as 15 (default setting).

### Data description

The 10x Visium human DLPFC dataset was collected from three independent adult samples. Each sample consists of four adjacent tissue slices. The manually annotated labels in the original study include white matter (WM) and six neocortical layers. We used these annotations as ground truth to evaluate the performance of simultaneous spatial domain identification and the quality of batch-corrected embeddings. Each slice contains ~4000 spots, ranging from 3498 to 4789. The mouse brain dataset was profiled by 10x Visium platform and the tissue was divided into two slices in the sagittal direction. The anterior and posterior slice contains 2,695 and 3355 spots, respectively. The adult mouse olfactory bulb dataset was profiled by Stereo-seq and Slide-seqV2 respectively. The Stereo-seq slice processed by an early study^9^ contains 19,109 spots. The Slide-seqV2 slice contains 20,139 spots. The early mouse embryo dataset consisting of four stages (E9.5, E10.5, E11.5, E12.5) profiled by Stereo-seq, contains 5913, 18408, 30124, 51365 spots, respectively. The ST data of normal and AD disease mouse hippocampus slices both profiled by Slide-seqV2 contain 19285 and 15092 spots, respectively. For the AD slice, an adjacent Aβ plaque staining image of the same brain tissue was also collected. The mouse hippocampus data consisting of seven adjacent slices profiled by Slide-seqV1 with the number of spots ranging from 14860 to 32895. Slide-seqV1 data are highly noisy and sparse with lower transcript detection sensitivity than Slide-seqV2^5^. Thus, recovering the true biology patterns is more challenging. Summary of these datasets can be found in **Supplementary Table 1**.

## Data availability

The datasets analyzed in this study are all from publicly available datasets. Specifically, the human DLPFC dataset can be accessed in the *spatialLIBD* package (http://spatial.libd.org/spatialLIBD). The mouse olfactory bulb tissue data generated by Stereo-seq and Slide-seqV2 platforms can be accessed from https://github.com/JinmiaoChenLab/SEDR_analyses, https://singlecell.broadinstitute.org/single_cell/study/SCP815, respectively. The mouse sagittal posterior and anterior brain data can be accessed at https://support.10xgenomics.com/spatial-gene-expression/datasets/1.0.0/V1_Mouse_Brain_Sagittal_Posterior; https://support.10xgenomics.com/spatial-gene-expression/datasets/1.0.0/V1_Mouse_Brain_Sagittal_Anterior. The mouse embryo can be accessed at https://db.cngb.org/stomics/mosta/. The normal and Alzheimer’s disease mouse hippocampus can be accessed at https://singlecell.broadinstitute.org/single_cell/study/SCP815 and https://singlecell.broadinstitute.org/single_cell/study/SCP1663, respectively. The Slide-seq 3D mouse hippocampus slice can be accessed at https://singlecell.broadinstitute.org/single_cell/study/SCP354/slide-seq-study. The annotation images from the Allen Mouse Brain Atlas can be accessed at https://mouse.brain-map.org/static/atlas (**Supplementary Table 2**).

## Acknowledgements

This work has been supported by the National Key Research and Development Program of China [Nos. 2021YFA1302500 and 2019YFA0709501 to S.Z.], the Strategic Priority Research Program of the Chinese Academy of Sciences [Nos. XDA16021400, XDPB17], the National Natural Science Foundation of China [Nos. 12126605, 61621003], the Key-Area Research and Development of Guangdong Province [No. 2020B1111190001], and the CAS Project for Young Scientists in Basic Research [No. YSBR-034 to S.Z.].

## Author contributions

S.Z. conceived and supervised the project. X.Z. developed and implemented the STAligner algorithm. X.Z., K.D. and S.Z. validated the methods and wrote the manuscript. All authors read and approved the final manuscript.

## References

1 Rao, A., Barkley, D., França, G. S. & Yanai, I. Exploring tissue architecture using spatial transcriptomics. Nature 596, 211–220 (2021).

2 Ståhl, P. L. et al. Visualization and analysis of gene expression in tissue sections by spatial transcriptomics. Science 353, 78–82 (2016).

3 Moses, L. & Pachter, L. Museum of spatial transcriptomics. Nature Methods 19, 534–546 (2022).

4 Rodriques, S. G. et al. Slide-seq: A scalable technology for measuring genome-wide expression at high spatial resolution. Science 363, 1463–1467 (2019).

5 Stickels, R. R. et al. Highly sensitive spatial transcriptomics at near-cellular resolution with Slide-seqV2. Nature biotechnology 39, 313–319 (2021).

6 Chen, A. et al. Spatiotemporal transcriptomic atlas of mouse organogenesis using DNA nanoball-patterned arrays. Cell 185, 1777–1792. e1721 (2022).

7 Maynard, K. R. et al. Transcriptome-scale spatial gene expression in the human dorsolateral prefrontal cortex. Nature neuroscience 24, 425–436 (2021).

8 Cable, D. M. et al. Cell type-specific inference of differential expression in spatial transcriptomics. Nature Methods, 1–12 (2022).

9 Fu, H. et al. Unsupervised spatially embedded deep representation of spatial transcriptomics. Preprint at https://www.biorxiv.org/content/10.1101/2021.06.15.448542v2. (2021).

10 Hu, J. et al. SpaGCN: Integrating gene expression, spatial location and histology to identify spatial domains and spatially variable genes by graph convolutional network. Nature methods 18, 1342–1351 (2021).

11 Zhao, E. et al. Spatial transcriptomics at subspot resolution with BayesSpace. Nature Biotechnology 39, 1375–1384 (2021).

12 Dong, K. & Zhang, S. Deciphering spatial domains from spatially resolved transcriptomics with an adaptive graph attention auto-encoder. Nature communications 13, 1–12 (2022).

13 Sun, S., Zhu, J. & Zhou, X. Statistical analysis of spatial expression patterns for spatially resolved transcriptomic studies. Nature methods 17, 193–200 (2020).

14 Zhang, C., Dong, K., Aihara, K., Chen, L. & Zhang, S. STAMarker: Determining spatial domain-specific variable genes with saliency maps in deep learning. Preprint at https://www.biorxiv.org/content/10.1101/2022.11.07.515535v1. (2022).

15 Shao, X. et al. Knowledge-graph-based cell-cell communication inference for spatially resolved transcriptomic data with SpaTalk. Nature Communications 13, 1–15 (2022).

16 Chen, S. et al. Spatially resolved transcriptomics reveals unique gene signatures associated with human temporal cortical architecture and Alzheimer’s pathology. Preprint at https://www.biorxiv.org/content/10.1101/2021.07.07.451554v1. (2021).

17 Korsunsky, I. et al. Fast, sensitive and accurate integration of single-cell data with Harmony. Nature methods 16, 1289–1296 (2019).

18 Stuart, T. et al. Comprehensive integration of single-cell data. Cell 177, 1888–1902. e1821 (2019).

19 Zeira, R., Land, M., Strzalkowski, A. & Raphael, B. Alignment and integration of spatial transcriptomics data. Nature Methods 19, 567–575 (2022).

20 Villacampa, E. G. et al. Genome-wide spatial expression profiling in formalin-fixed tissues. Cell Genomics 1, 100065 (2021).

21 Yao, Z. et al. A taxonomy of transcriptomic cell types across the isocortex and hippocampal formation. Cell 184, 3222–3241. e3226 (2021).

22 Kadowaki, K. et al. Phosphohippolin expression in the rat central nervous system. Molecular brain research 125, 105–112 (2004).

23 Zacharias, D. A. & Kappen, C. Developmental expression of the four plasma membrane calcium ATPase (Pmca) genes in the mouse. Biochimica et Biophysica Acta-General Subjects 1428, 397–405 (1999).

24 Mamoor, S. The α1 subunit of the γ-aminobutyric acid receptor,control the limited hepatocyte Gabra1, is differentially expressed in the brains of patients with schizophrenia. Preprint at https://doi.org/10.31219/osf.io/m93ya. (2020).

25 Richardson, L. et al. EMAGE mouse embryo spatial gene expression database: 2014 update. Nucleic acids research 42, D835–D844 (2014).

26 Savolainen, S. M., Foley, J. F. & Elmore, S. A. Histology atlas of the developing mouse heart with emphasis on E11. 5 to E18. 5. Toxicologic pathology 37, 395–414 (2009).

27 Haddad-Tóvolli, R., Szabó, N.-E., Zhou, X. & Alvarez-Bolado, G. Genetic manipulation of the mouse developing hypothalamus through in utero electroporation. JoVE (Journal of Visualized Experiments), e50412 (2013).

28 DíDz-Guerra, E., Pignatelli, J., Nieto-Estrra, E., Pignatell-Abejra, E., Pignatelli, J., Nietoiments) X. & Alvarez-Bolado, G. Genet The Anatomical Record 296, 1364–1382 (2013).

29 Chen, V. S. et al. Histology atlas of the developing prenatal and postnatal mouse central nervous system, with emphasis on prenatal days E7. 5 to E18. 5. Toxicologic pathology 45, 705–744 (2017).

30 Siddiqui, T. J. et al. An LRRTM4-HSPG complex mediates excitatory synapse development on dentate gyrus granule cells. Neuron 79, 680–695 (2013).

31 Elmore, M. R. et al. Colony-stimulating factor 1 receptor signaling is necessary for microglia viability, unmasking a microglia progenitor cell in the adult brain. Neuron 82, 380–397 (2014).

32 Zhang, B. et al. Integrated systems approach identifies genetic nodes and networks in late-onset Alzheimer’s disease. Cell 153, 707–720 (2013).

33 Zhao, Y. et al. TREM2 is a receptor for β-amyloid that mediates microglial function. Neuron 97, 1023–1031. e1027 (2018).

34 Hong, S. et al. Complement and microglia mediate early synapse loss in Alzheimer mouse models. Science 352, 712–716 (2016).

35 Zeisel, A. et al. Molecular architecture of the mouse nervous system. Cell 174, 999–1014. e1022 (2018).

36 Tzingounis, A. V., Kobayashi, M., Takamatsu, K. & Nicoll, R. A. Hippocalcin gates the calcium activation of the slow afterhyperpolarization in hippocampal pyramidal cells. Neuron 53, 487–493 (2007).

37 Yan, C., Costa, R., Darnell Jr, J. E., Chen, J. & Van Dyke, T. Distinct positive and negative elements control the limited hepatocyte and choroid plexus expression of transthyretin in transgenic mice. The EMBO Journal 9, 869–878 (1990).

38 Rood, J. E. et al. Toward a common coordinate framework for the human body. Cell 179, 1455–1467 (2019).

39 Wolf, F. A., Angerer, P. & Theis, F. J. SCANPY: large-scale single-cell gene expression data analysis. Genome biology 19, 1–5 (2018).

40 Salehi, A. & Davulcu, H. Graph attention auto-encoders. arXiv preprint arXiv:.10715 (2019).

41 Simon, L. M., Wang, Y.-Y. & Zhao, Z. Integration of millions of transcriptomes using batch-aware triplet neural networks. Nature Machine Intelligence 3, 705–715 (2021).

42 Dong, Z. et al. Registration of large-scale terrestrial laser scanner point clouds: A review and benchmark. ISPRS Journal of Photogrammetry Remote Sensing 163, 327–342 (2020).

43 Umeyama, S. Least-squares estimation of transformation parameters between two point patterns. IEEE Transactions on Pattern Analysis Machine Intelligence 13, 376–380 (1991).

44 Kingma, D. P. & Ba, J. Adam: A method for stochastic optimization. arXiv preprint arXiv:. (2014).

45 Clevert, D.-A., Unterthiner, T. & Hochreiter, S. Fast and accurate deep network learning by exponential linear units (elus). Preprint at https://arxiv.org/abs/1511.07289. (2015).

46 Fey, M. & Lenssen, J. E. Fast graph representation learning with PyTorch Geometric. Preprint at https://arxiv.org/abs/1903.02428. (2019).

47 Fraley, C., Raftery, A. E., Murphy, T. B. & Scrucca, L. mclust version 4 for R: normal mixture modeling for model-based clustering, classification, and density estimation. (Technical report, 2012).

48 Yu, G., Wang, L.-G., Han, Y. & He, Q.-Y. clusterProfiler: an R package for comparing biological themes among gene clusters. Omics: a journal of integrative biology 16, 284–287 (2012).

49 Tran, H. T. N.et al. A benchmark of batch-effect correction methods for single-cell RNA sequencing data. Genome biology 21, 1–32 (2020).

