## Supplementary Text, Figures and Tables. for "Integrating spatial transcriptomics data across different conditions, technologies, and developmental stages"

### Supplementary Notes

#### Evaluation metrics for comparison of integration methods.

We employed the divergence score to quantify the performance in mixing the shared spatial domains across slices after integration. A smaller divergence score means more sufficient mixing of the same spatial domain. The degree of mixing is measured by the universal *k*-nearest-neighbor (kNN) divergence[1]. For the shared domain $l$ across two slices with corresponding UMAP embeddings $h_{1,l}$ and $h_{2,l}$, we define the kNN divergence as:

$$D_{kNN}\left(h_{1,l}||h_{2,l}\right) = \frac{d}{n}\sum_{i=1}^{n} \log\frac{v(i)}{\rho(i)} + \log\frac{m}{n-1}, \quad (1)$$

where $d$ is the size of spot embedding, $m$ and $n$ are the numbers of spots in domain $l$ in slices 1 and 2, respectively, $\rho(i)$ is the Euclidean distance between spot $i$ and its kNNs in the same slice, and $v(i)$ is the distance from spot $i$ to its kNNs in the other slice. The divergence score is defined as the average kNN divergence over all spatial domains. We set $k$ as 30 for the kNN computation.

We further evaluated the separation of different spatial domains by the silhouette coefficient which measures the average distance between any two domains. A higher silhouette score means better domain separation after batch correction. We denote $a(i)$ as the average distance between spot $i$ to all the other spots in the same domain and $b(i)$ as the average distance between spot $i$ and spots in the next closest domain. The distance is computed in the UMAP embedding space of the batch-corrected data. Here, the Euclidean distance is used. The silhouette coefficient of spot $i$ is written as

$$S(i) = \frac{b(i)-a(i)}{\max\{a(i),b(j)\}} \in [-1, 1], i = 1, \dots, n. \quad (2)$$

The average silhouette coefficient across all spots is computed as the silhouette score.

### Figures

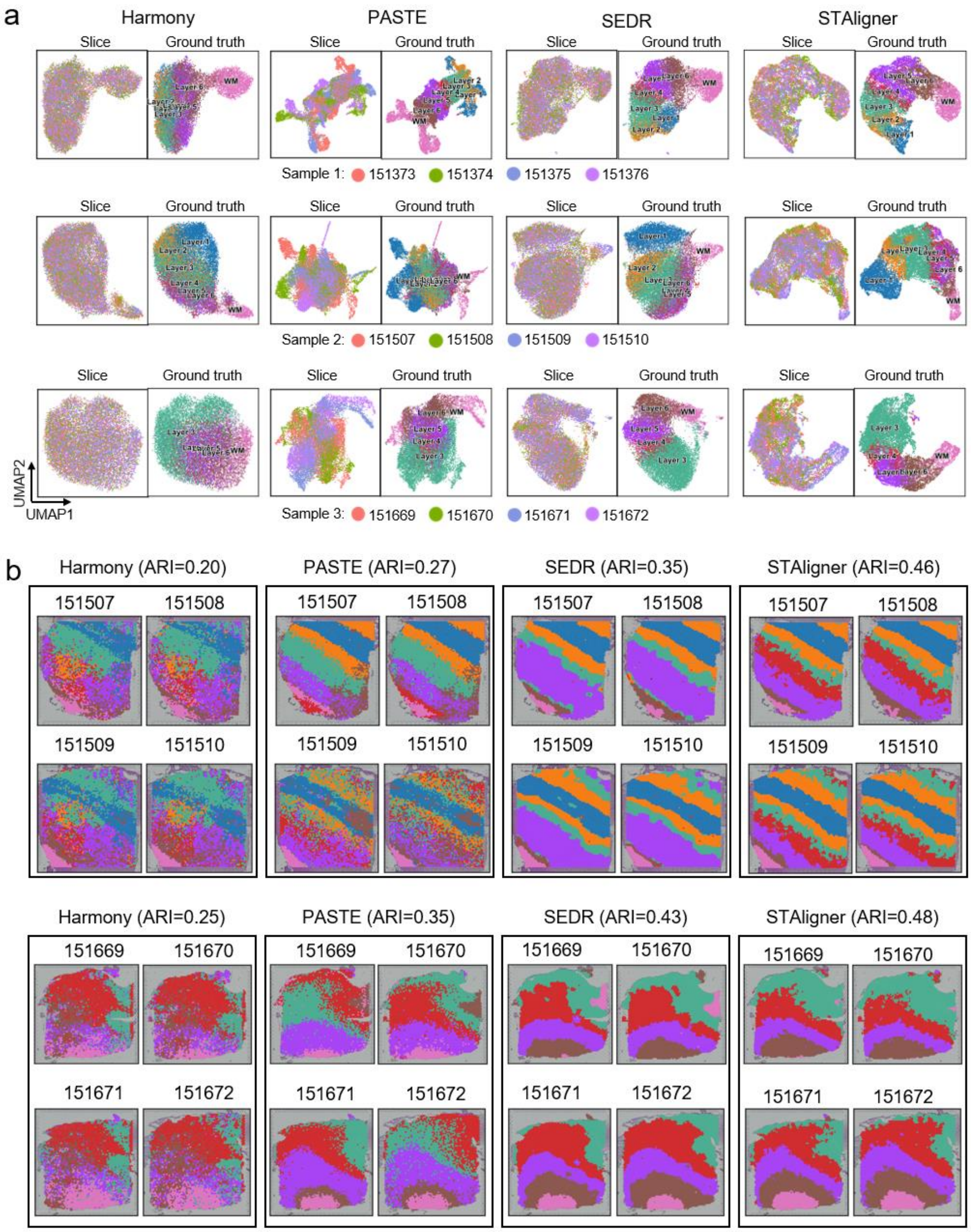


**Fig. S1.** Integration of the four slices of samples 1, 2 and 3 by different methods respectively, related to **Fig. 2**. **a.** UMAP plots of embeddings colored by slices (left) and cortical layers (right) for the four integration methods in samples 1, 2 and 3 respectively. **b.** Spatial domains of the consecutive slices identified simultaneously in samples 2 and 3 by the four integration methods, respectively.

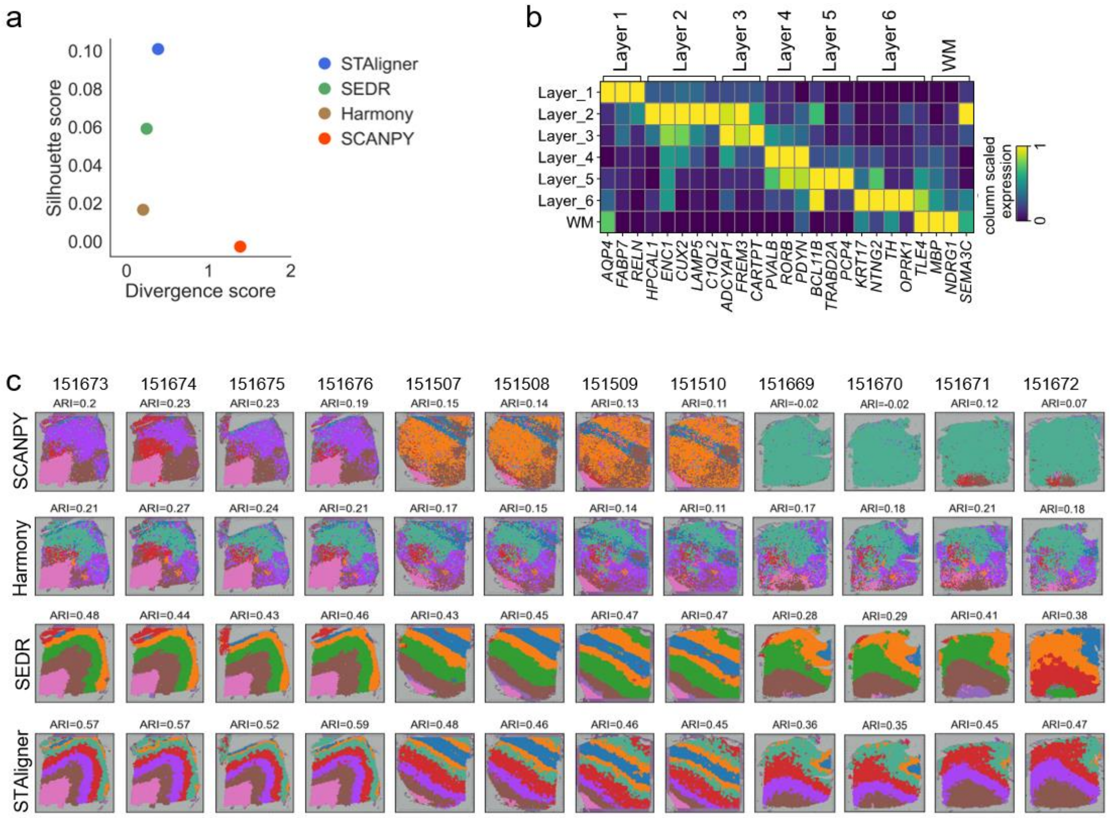


**Fig. S2.** Integration of all the 12 slices of samples 1, 2 and 3, related to **Fig 2**. **a.** Scatter diagram of the integration quality measured by the divergence score versus the silhouette score. **b.** Mean expressions of the corresponding marker genes across the cortical layers identified by STAligner. The expressions were scaled by genes. **c.** Spatial domains of each slice identified simultaneously across all slices by the four integration methods, respectively.

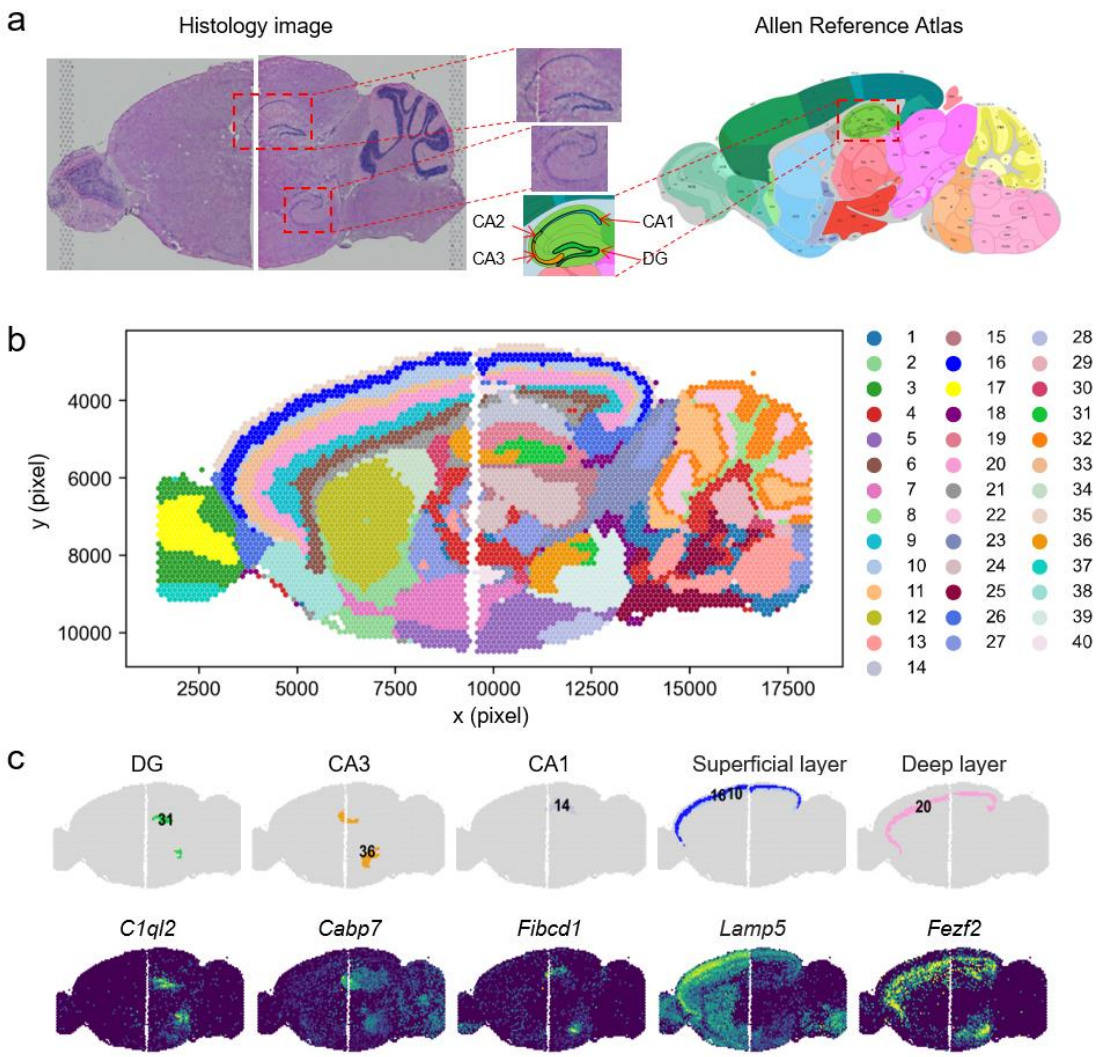


**Fig. S3.** STAligner can stitch adjacent mouse brain sagittal anterior and posterior slices to generate a large composite slice. **a.** The H&E images of each slice and the corresponding anatomical definitions obtained from the Allen Mouse Brain Atlas. **b.** Visualization of spatial domains identified simultaneously across the two slices by STAligner with mclust clustering. **c.** The tissue structures annotated based on the Allen Mouse Brain Atlas and their corresponding marker genes.

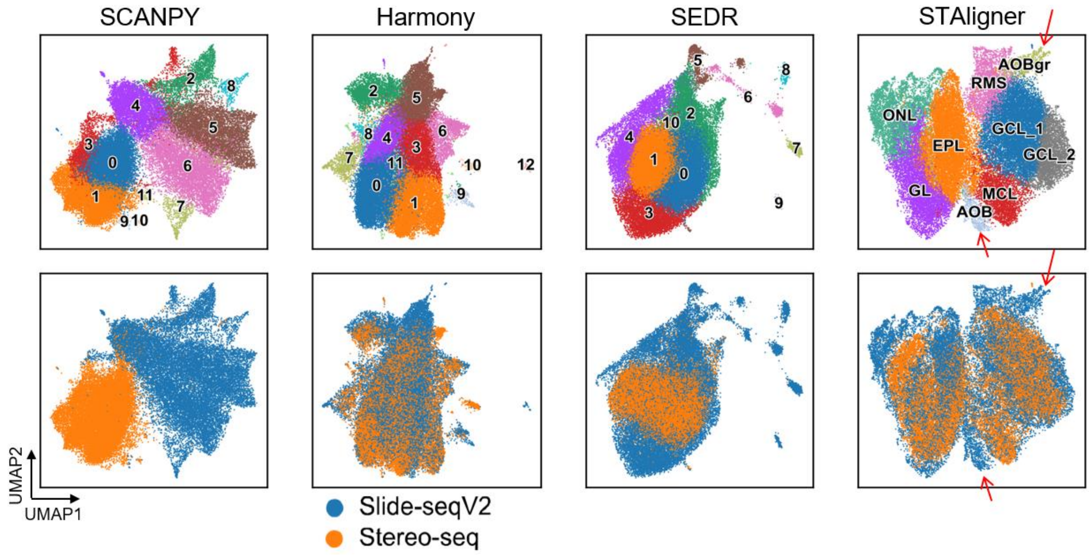


**Fig. S4.** UMAP plots of embeddings colored by spatial domains (top) and slices (bottom) for SCANPY, Harmony, SEDR and STAligner respectively, related to **Fig 3**.

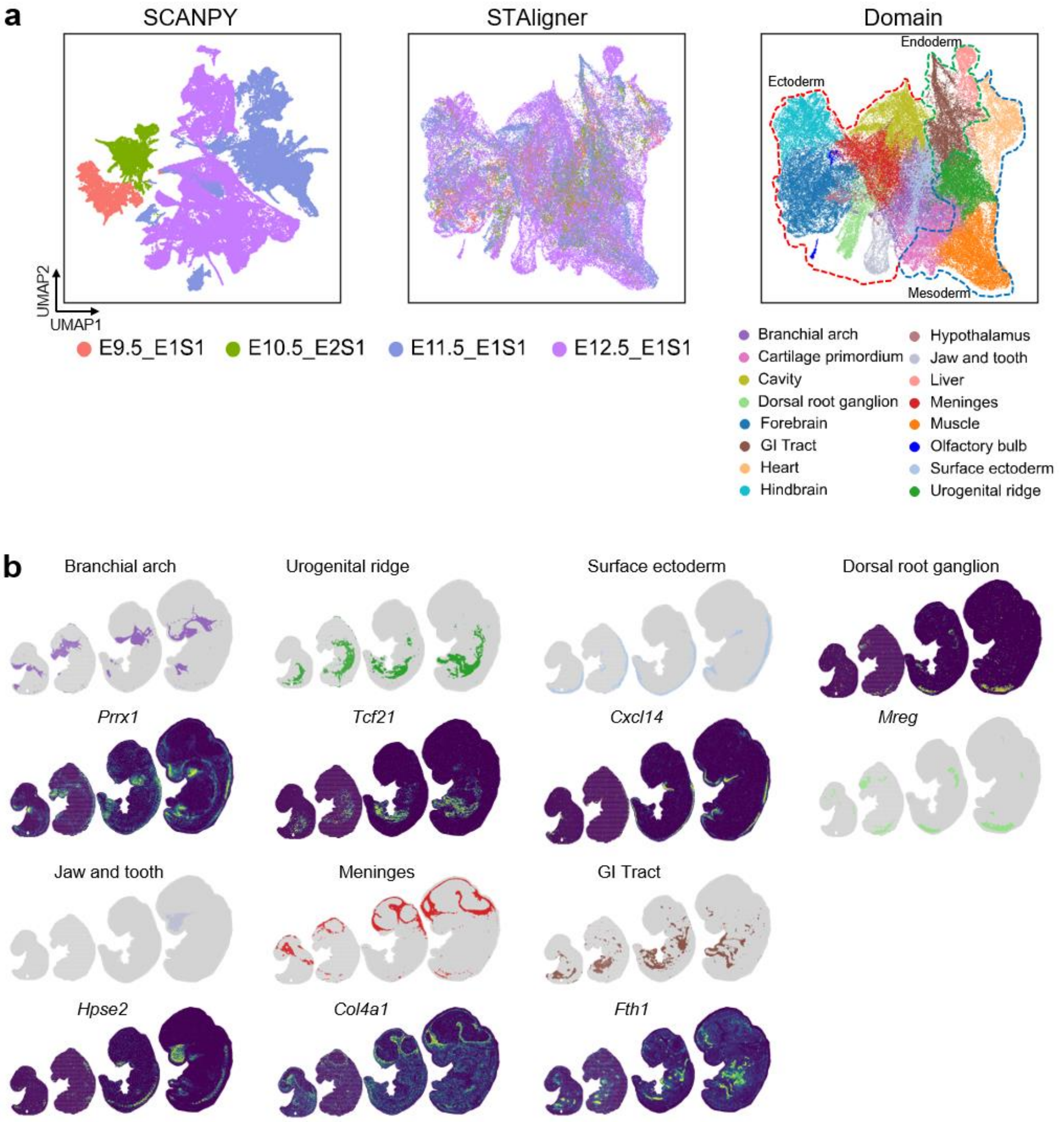


**Fig. S5.** Integration of early developmental ST slices of the mouse embryo produced by Stereo-seq, related to **Fig 4**. **a.** UMAP plots of SCANPY and STAligner embeddings colored by developmental stages and spatial domains. **b.** Spatial tissue substructures relating to branchial arch, urogenital ridge, surface ectoderm, dorsal root ganglion, jaw and tooth, meninges and GI tract (top) and their corresponding marker genes (bottom).

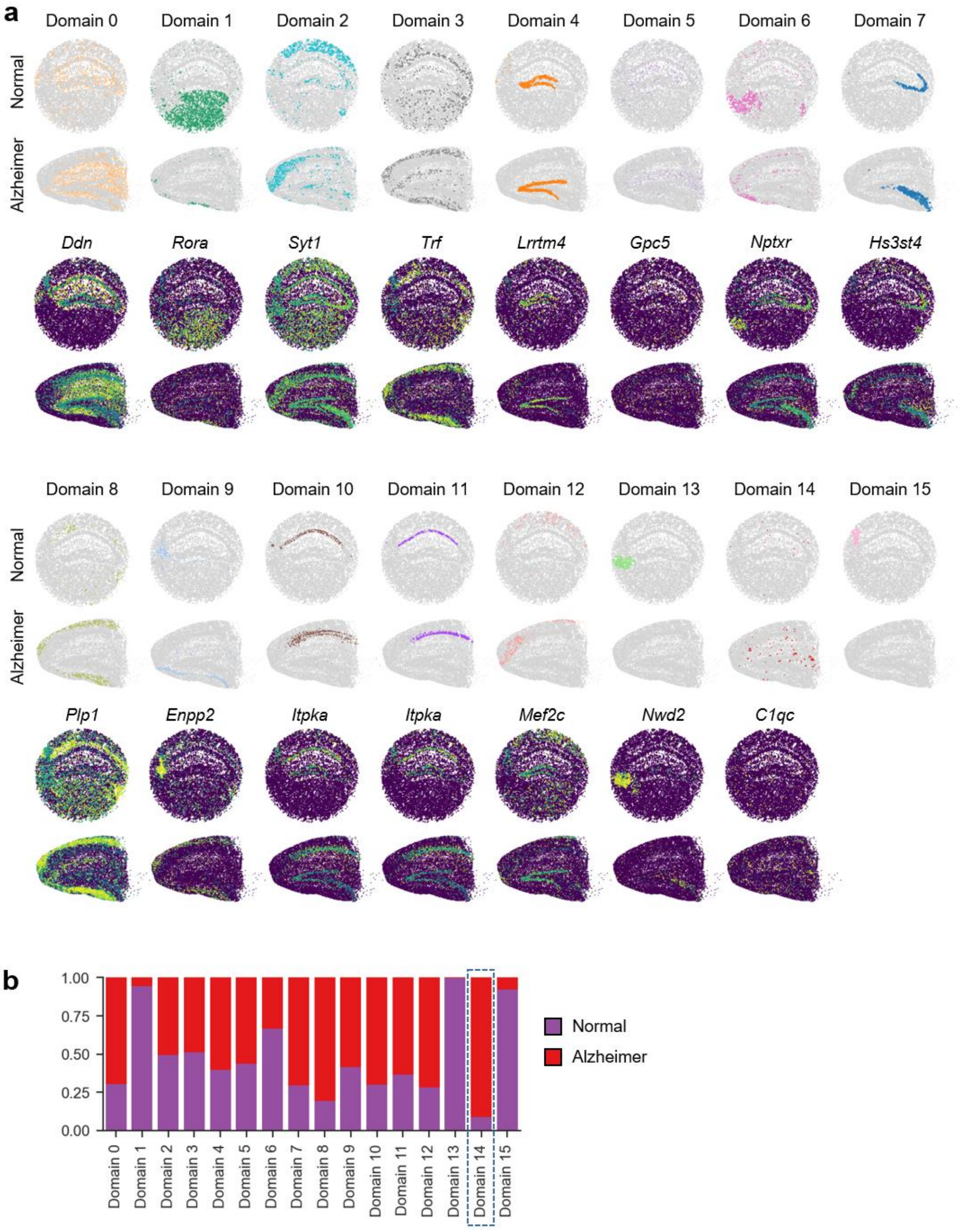


**Fig. S6.** Integration of two mouse hippocampus slices of normal and Alzheimer's disease conditions profiled by Slide-seqV2, related to **Fig 5**. **a.** Spatial visualization of spatial domains identified simultaneously across two slices by STAligner (top) and the expressions of their corresponding marker genes (bottom). **b.** Bar plot showing the proportion of spots of each domain belonging to normal and Alzheimer's slices to identify disease-specific domain (dashed box).

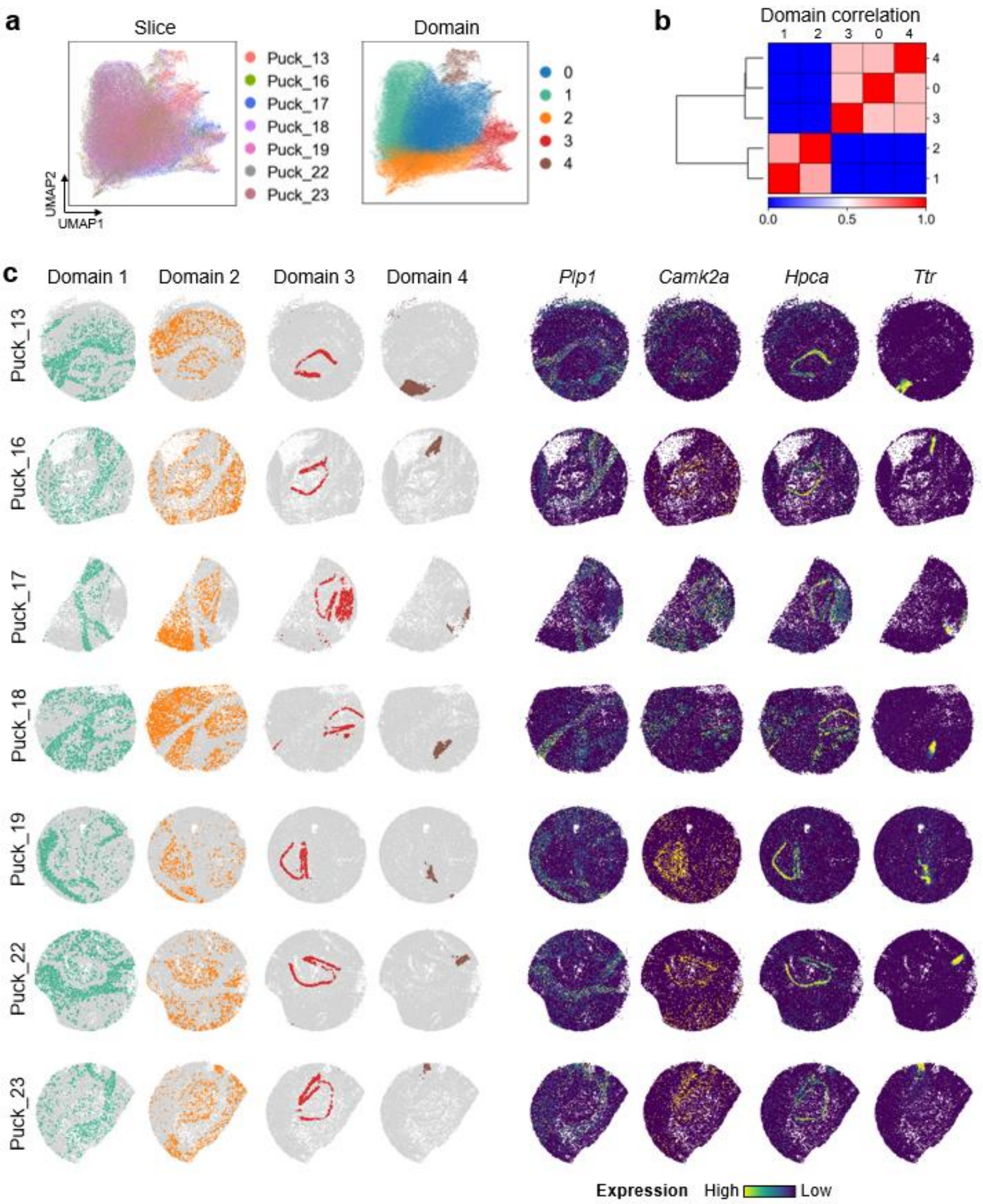


**Fig. S7.** Integration of seven adjacent mouse hippocampus slices profiled by Slide-seq, related to **Fig 6**. **a.** UMAP plots of STAligner embeddings colored by the seven adjacent slices (left) and spatial domains (right), respectively. **b.** Pearson correlation coefficient matrix of the identified spatial domains. **c.** Spatial visualization of spatial domains 1, 2, and 4 identified simultaneously across all slices by STAligner and the expressions of their corresponding marker genes.

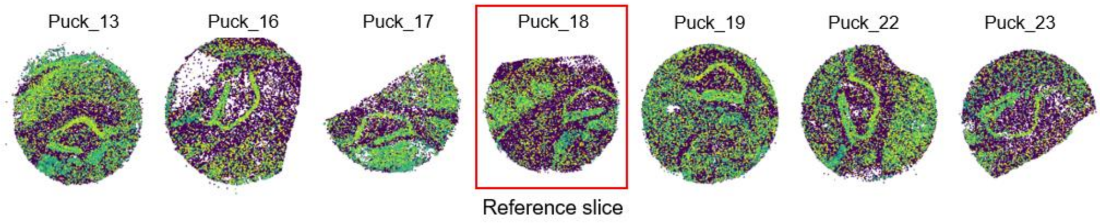


**Fig. S8.** Pairwise alignment of slices by PASTE, related to **Fig 6**. Puck_18 was considered as the reference slice and the spatial coordinates of the other slices are projected to the spatial coordinate system of Puck_18.

**a**

| Tissue | # of slices | # of spots | Time | Memory |
|---|---|---|---|---|
| Human dorsolateral prefrontal cortex (DLPFC) | 12 | 47329 | 116 s | 4.3 GB |
| Mouse embryos | 4 | 95896 | 250 s | 12.8 GB |
| Mouse hippocampus | 7 | 171645 | 503 s | 15.6 GB |

**b**

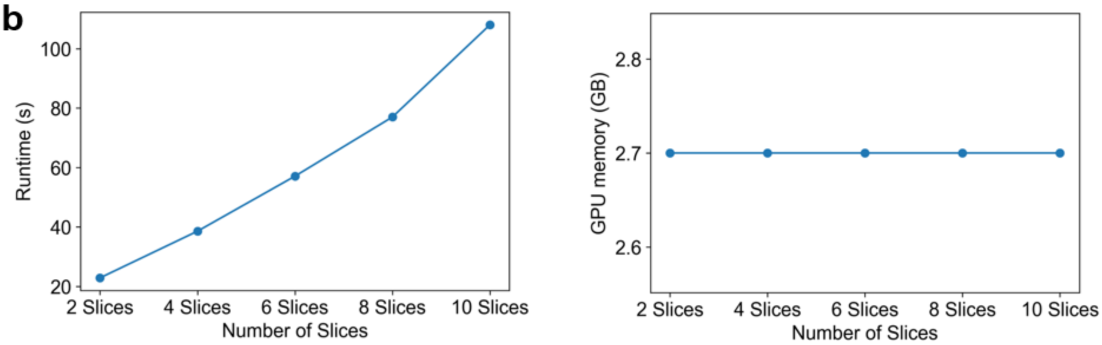


**Fig. S9.** Computational time and GPU memory usage cost by STAligner on three real datasets (**a**) and the simulated dataset (**b**) respectively. We generated the simulated dataset by replicating the number of the DLPFC slice 151673 and evaluated the scalability of STAligner. The experiments were performed on a GPU server with Intel(R) Core(TM) i9-10920X CPU @ 3.50GHz and NVIDIA GeForce RTX 3090 GPU.

**Table S1. Summary of the ST data used in this study.**

| Tissue | Platform | Slice id | # of spots | Related figures |
|---|---|---|---|---|
| **Human dorsolateral prefrontal cortex (DLPFC)** | 10x Visium | 151507, 151508, 151509, 151510, 151669, 151670, 151671, 151672, 151673, 151674, 151675, 151676. | 4226, 4384, 4789, 4634, 3661, 3498, 4110, 4015, 3639, 3673, 3592, 3460. | Fig. 2 Fig. S1, S2 |
| **Mouse brain** | 10x Visium | Sagittal-anterior | 2695 | Fig. S3 |
| | | Sagittal-posterior | 3355 | |
| **Mouse olfactory bulb** | Slide-seqV2 | Puch_180531_23 | 20139 | Fig. 3 |
| | Stereo-seq | N.A. | 19109 | Fig. S4 |
| **Mouse embryos** | Stereo-seq | E9.5_E1S1, E10.5_E2S1, E11.5_E1S1, E12.5_E1S1. | 5913, 8494, 30124, 51365. | Fig. 4 Fig. S5 |
| **Mouse hippocampus** | Slide-seqV2 | Alzheimer's j20_rep1 | 15092 | Fig. 5 Fig. S6 |
| | | Normal Puck_190921_21 | 12822 | |
| **Mouse hippocampus** | Slide-seqV1 | Puck_180531_13, Puck_180531_16, Puck_180531_17, Puck_180531_18, Puck_180531_19, Puck_180531_22, Puck_180531_23. | 32895, 18631, 14860, 26912, 31163, 28676, 18508. | Fig. 6 Fig. S7, S8 |

**Table S2. The slice ID and URL to get the anatomic reference data from the Allen Mouse Brain Atlas.**

| Related figures | ABA ID | ABA URL |
|---|---|---|
| Fig. S3a | 100883818 | http://atlas.brain-map.org/atlas?atlas=2&plate=100883818 |
| Fig. 3b | 100960460 | http://atlas.brain-map.org/atlas?atlas=1&plate=100960460 |
| Fig. 5a | 100960084 | http://atlas.brain-map.org/atlas?atlas=1&plate=100960084 |

### Reference

1 Wang, Q., Kulkarni, S. R. & Verdú, S. Divergence estimation for multidimensional densities via *k*-Nearest-Neighbor distances. *IEEE Transactions on Information Theory* **55**, 2392-2405 (2009).
